# The influence of everyday events on prospective timing “in the moment”

**DOI:** 10.1101/246975

**Authors:** Ashley S. Bangert, Christopher A. Kurby, Jeffrey M. Zacks

**Author notes:** Corresponding Author: Ashley S. Bangert, Address: Department of Psychology, The University of Texas at El Paso, 500 W. University, Ave., El Paso, Texas, 79902.

## Abstract

We conducted two experiments to investigate how the eventfulness of everyday experiences influences people’s prospective timing ability. Specifically, we investigated whether events contained within movies of everyday activities serve as markers of time, as predicted by Event Segmentation Theory, or whether events pull attention away from the primary timing task, as predicted by the Attentional Gate theory. In the two experiments reported here, we asked participants to reproduce a previously learned 30 second target duration while watching a movie that contained eventful and uneventful intervals. In Experiment 2, reproduction also occurred during “blank movies” while watching a fixation. In both experiments, participants made shorter and more variable reproductions while simultaneously watching eventful as compared to uneventful movie intervals. Moreover, in Experiment 2, the longest reproductions were produced when participants had to watch the blank movies, which contained no events. These results support Event Segmentation Theory and demonstrate that the elapsing events during prospective temporal reproduction appear to serve as markers of temporal duration rather than distracting from the timing task.

## Introduction

Keeping track of time is ubiquitous in many everyday activities. For example, a caterer must monitor elapsing time during a party to know when to serve particular dishes to keep things on schedule. After the party is over the caterer may try to remember how long it took to serve the main course to better plan for her next event. If the caterer did not have a clock available to keep track of time, how might she make her temporal judgments?

People rely on different processes in situations where they know in advance they need to keep track of time (*prospective timing*) versus when they must estimate, after the fact, how long a prior experience lasted (*retrospective timing*) (see Block & Gruber, 2014; Block & Zakay, 1997 for reviews). For retrospective timing, people use their memory for the number and coherence of contextual changes to make temporal estimates; estimates increase with more changes or when changes are unpredictable or incoherent (Block & Reed; 1978; Boltz, 1995; Poynter, 1983; Zauberman, Levav, Diehl & Bhargrave, 2010). Under prospective timing conditions, the degree to which attentional resources are allocated to timing is crucial; duration estimates decrease when attention is divided between timing and a concurrent task (Brown, 1997; Macar, Grondin & Casini, 1994).

Attentional Gate Theory (AGT) (Zakay & Block, 1995) is typically used to explain this phenomenon in prospective timing. When a stimulus is encountered, pulses from an internal pacemaker pass through an attentional gate to an accumulator that temporarily stores the total pulses as a representation of the current temporal interval. The more attention paid to time, the wider the gate opens. If attention is allocated, instead, to non-temporal task features, the gate is narrowed and pulses are missed. Thus, if someone pays attention to non-temporal information while reproducing a previously encoded interval, the reproduction will be longer than the initially encoded duration. Consider the caterer: If she attends to non-temporal features of the catering event she will miss pulses, and time will seem to pass more slowly.

However, there is reason to think that the structure of activity may contribute to the prospective perception of the passage of time. People tend to segment the stream of sensory and contextual information in their everyday experience into meaningful spatio-temporal units, known as events (Newtson, 1976; Zacks, Speer, Swallow, Braver & Reynolds, 2007). Event Segmentation Theory (EST; Kurby & Zacks, 2008; Zacks et al., 2007) argues that people create event models, which are working memory representations of what is currently happening, to help them predict what might happen next. At event boundaries, prediction errors increase and the event model is updated. Many perceptual or semantic cues can signal an event change, including a change in the goal of the actor, a change in spatial location, etc. (Zacks, Speer & Reynolds, 2009). The perception of event boundaries confers many processing consequences--working memory is updated (Bailey, Kurby, Sargent, & Zacks, 2017), visual attention is modulated (Eisenberg & Zacks, 2016), and new episodic memories are encoded (Ben-Yakov, Dudai, & Eshai, 2013; Ezzyat & Davachi, 2011). Given this, one might expect that the number of events during an ongoing experience will influence people’s prospective estimates of time.

Indeed, recent work suggests that, similar to retrospective timing, the number and quality of contextual changes and event structure may influence prospective timing beyond the traditional effects of attention. For example, simple naturalistic stimuli that violate expectations lead to longer prospective judgments than predictable stimuli (Boltz, 2005). Some studies have also found that more contextual changes or segments lead people to think more time has passed. For example, Faber & Gennari (2017) found that when asked to attend to time, the more segments participants perceived in unfamiliar animations, the longer their subsequent duration estimates. Waldum and Sahakyan (2013) further discovered that people used their knowledge of the number and typical length of pop songs to make prospective estimates of the length of time they spent completing a lexical decision task (LDT) with background music or to complete a time-based prospective memory (TBPM) task during the LDT. The more songs played, the longer they judged the LDT to be and the earlier they thought a 10 minute TBPM target had passed. However, some studies have found the opposite result. Liverence and Scholl (2012), for example, found that stimuli with more perceived segments were judged as shorter (see also Predebon, 1996; Sargent, Zacks, Philbeck, & Flores, 2013). Thus, it is still unclear how and when contextual changes exert their influence on prospective timing. Many studies demonstrating the typical dissociation between processes involved in prospective and retrospective timing have used artificial stimuli; timing under naturalistic experiences where temporal and non-temporal features are tightly bound together may show a stronger influence of non-temporal contextual influences.

Most prior studies that showed contextual change or segmentation effects on prospective timing manipulated segmentation at encoding and asked participants to later make estimates or reproductions of these encoded intervals. However, in naturalistic prospective timing experiences, individuals monitor time concurrently with the unfolding of events. The caterer, for example, must monitor time as dishes are served, food is periodically replenished, dishes are collected, etc. Here, we assessed the effects of the passing of events when concurrently engaging in prospective temporal reproductions. In two experiments, participants learned a temporal duration then reproduced that duration while watching movies of actors engaged in everyday activities. If, as suggested by AGT, the processing of events pulls attention away from timing, participants should make longer reproductions during eventful than uneventful movie intervals, because it will take longer to accumulate the pulses that match the previously encoded target. Alternatively, if event boundaries are contextual changes that serve as markers of time, consistent with EST, then participants should make shorter reproductions during eventful than uneventful intervals because they will perceive time as moving more quickly.

### Experiment 1

Participants reproduced a target duration while watching eventful or uneventful intervals of movies. Eventful intervals were windows of time that contained many event boundaries, and uneventful intervals contained few event boundaries. Prior to watching each movie, participants encoded the target duration while watching a fixation.

## Method

### Participants

Fifty college students with normal vision and hearing completed the study for course credit. The experiment was approved by the Washington University Institutional Review Board. We eliminated five participants who failed to make reproductions on 33% or more of the trials and three who used a counting strategy. The remaining forty-two participants (mean age = 19.67 ± 1.84, 24 females) were included in the analyses.

### Apparatus and Stimuli

The experiment was administered using E-Prime software on Dell desktop computers. Participants were seated approximately 54 cm from the computer screen and were cued while watching movies of everyday activities to reproduce a target 30 second interval on which they had previously been trained. The reproduction cue was a 1000 Hz, 50 ms square tone. Movies were presented in a window positioned in the center of the screen that subtended approximately 21.7° (width) x 15.4° (height) of visual angle. The movies showed people making a sandwich, washing a car, doing laundry, pitching a tent, planting a flower box, and building a ship out of Duplo blocks. Figure 1 shows screen shots from each movie. The sandwich video was used during practice and the other five movies were used during experimental trials. Twelve eventful and twelve uneventful movie intervals were used for the experimental trials. Table 1 shows the movie lengths as well as the number of intervals identified from each movie.

**Figure 1.**
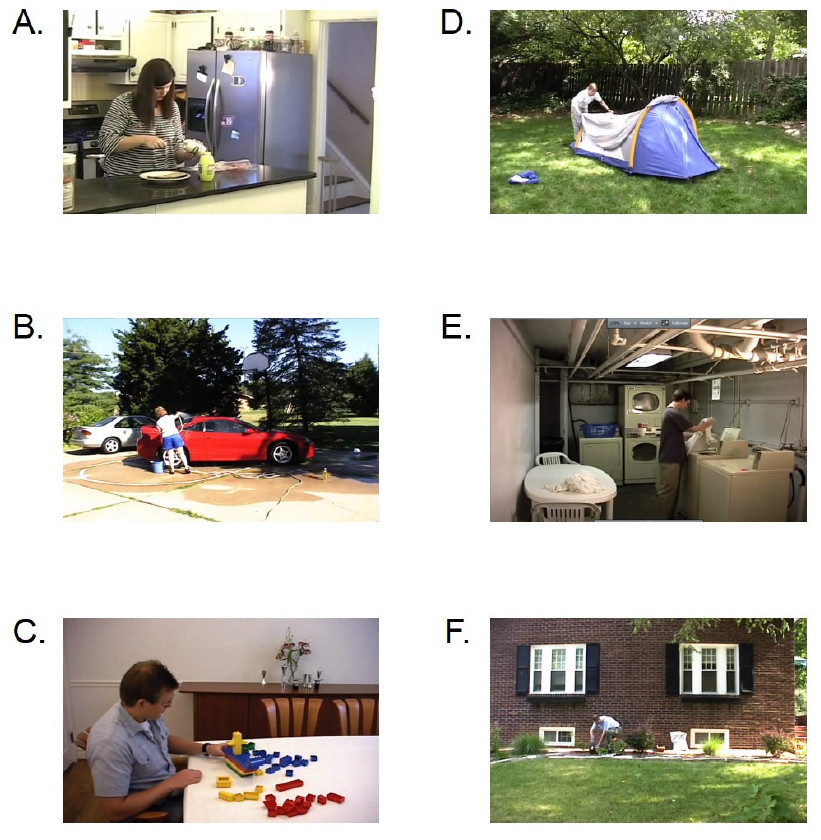
Screenshots from the everyday movies used in experiments 1 and 2. Panel A shows an image from the sandwich movie used for practice trials. Panel B shows an image from the washing a car movie. Panel C shows an image from the movie of a man building a ship out of Duplo blocks. Panel D shows an image of a woman pitching a tent. Panel E shows an image of the washing clothes movie. Panel F shows an image of the movie of a man planting a flower box.

**Table 1.** Experimental Movie Details from Experiments 1 and 2

| Label | Description | Length (sec) | # EV<br>Intervals | # UN<br>Intervals |
| --- | --- | --- | --- | --- |
| A | Washing a car | 432 | 3 | 2 |
| B | Building a Duplo<br>blocks ship | 246 | 2 | 2 |
| C | Pitching a tent | 379 | 2 | 3 |
| D | Washing clothes | 300 | 2 | 2 |
| E | Planting a flower box | 354 | 3 | 3 |
| Blank 1 | Blank Movie | 396 | - | - |
| Blank 2 | Blank Movie | 331 | - | - |

*Note.* EV refers to eventful intervals. UN refers to uneventful intervals. The length of the blank movies from experiment 2 are also displayed. There were no eventful or uneventful intervals for blank movies.

Movie intervals were selected using used fine-grained segmentation data obtained from younger and older adults in a previous study (Kurby & Zacks, 2011), in which participants pressed a button to indicate when they perceived an event boundary, defined as “when one meaningful unit of activity ended and another began.” From the data for each movie, we defined a 30 s moving window, starting at the onset of the movie, and counted the number of boundaries identified. That count was logged, the window was moved over one second, the next count was produced, and so on. The resulting total number of event boundaries per window was converted to a z-score which represented that window’s level of eventfulness. Then, all z-scores for the 30 s windows were rank ordered within movie. Starting with a randomly selected movie we first selected the most eventful 30 s time interval. That interval, the following 20 s, and any remaining intervals that overlapped this 50-s interval were then removed from the pool of windows. We then repeated this process, selecting the *least* eventful interval from what remained. This was repeated, alternating between eventful and uneventful selections until we selected 2 to 3 of each interval type for each movie, or ran out of available windows. The average z-score for eventful intervals was 1.61 with a range of 3.76; the average for uneventful intervals was -1.58 with a range of 1.80.

### Procedure

Participants completed the practice movie followed by the other 5 movies. Movie order was counterbalanced across participants. Participants were told to pay attention to time and to what was happening in each movie for a later order memory task. They were also instructed not to count. At the end of the experiment, participants completed an exit questionnaire asking about any strategies they used.

Prior to each movie, participants were trained on a 30 s target interval (see Figure 2). In two exposure trials, a green square appeared paired with a brief 1000 Hz 50 ms tone. Participants had to attend to how long the square remained until it turned red. Participants then reproduced the target interval during two baseline trials with visual feedback. During a baseline trial, a 1000 Hz 50 ms tone signaled participants to begin their reproduction and to press the spacebar once they thought the target duration had passed. The feedback display showed a black bar representing the target duration length above a red bar representing the participant’s reproduction. If, for example, their reproduced duration was 6% longer than the target, the red bar was 6% longer than the black bar. After training, participants reproduced the target interval during the movie. Reproduction cues were presented at the beginning of 30 s eventful or uneventful movie intervals and participants pressed the spacebar to end each reproduction.

**Figure 2.**
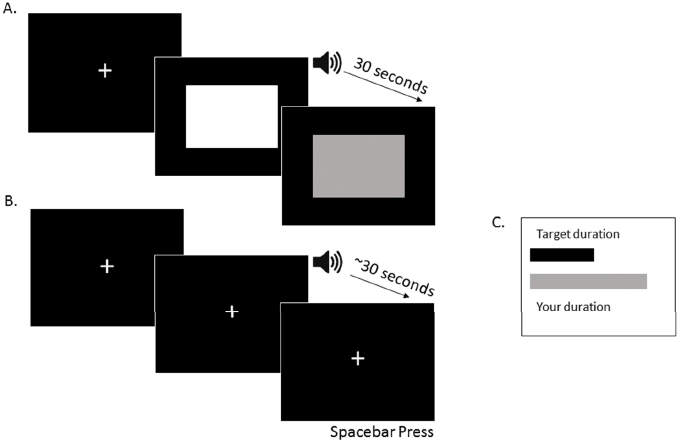
Schematic showing training trials for Experiments 1 and 2. Panel A shows the initial exposure training trials where participants first encoded the target interval. In the experiment, the white square was green and the grey square was red. Panel B shows the baseline trials where participants reproduced the duration with a spacebar press after hearing the tone which cued them to begin their reproduction. Panel C shows the feedback screen participants saw after each baseline trial demonstrating the relative length of their reproduction compared to the target interval. The gray bar representing the duration the participant reproduced was red during the actual experiment. The example here shows a baseline trial where the participant made a longer reproduction than the target interval.

After each movie, participants completed the order memory task where they reordered a set of scrambled 3 x 5 printed screenshots from the movie (6 images for practice; 12 images for each experimental movie) presented in two equal rows on a table. They had to reorder the images according to the order in which the presented activities occurred in the movie. We recorded total time to complete the task and the mean absolute deviation of each card’s sorted numeric position from the correct numeric position (Zacks et al., 2006).

### Data Analysis

We eliminated reproductions that were ± 3 standard deviations around the group mean across all 24 movie intervals. The same trimming procedure was applied to baseline trials. This eliminated an average of .31 experimental trials and .12 baseline trials per participant. We calculated the accuracy index (AI) for each remaining trial, which is the reproduced duration divided by the target duration. A value of 1 indicates a perfect reproduction, while values less than 1 and greater than 1 represent reproductions shorter and longer than the target duration, respectively. For each participant, we calculated the mean AI for baseline trials, eventful reproductions, and uneventful reproductions. The coefficient of variation (CV; standard deviation/mean reproduced duration) was also calculated for eventful and uneventful trials within each participant. CV is a variability measure that accounts for individual differences in mean reproductions. We conducted paired samples t-tests to compare AI and CV scores for reproductions made during eventful and uneventful movie intervals.

## Results

### Order Memory Task

The average completion time was 105.36 seconds (SD = 30.14) and the average deviation score was .54 (SD = .34), which shows that, on average, participants sorted images from the movies with high accuracy, suggesting they attended to the movie content.

### Accuracy Index (AI)

For baseline trials, participants achieved *M* = 1.03, which was less than one second longer than the actual 30 s target duration, suggesting they were trained appropriately. Importantly, there was a significant difference in AI for uneventful versus eventful movie intervals, *t*(41) = 2.47, *p* = .018, *d* = .21, with people making shorter reproductions during eventful trials (see Figure 3).

**Figure 3.**
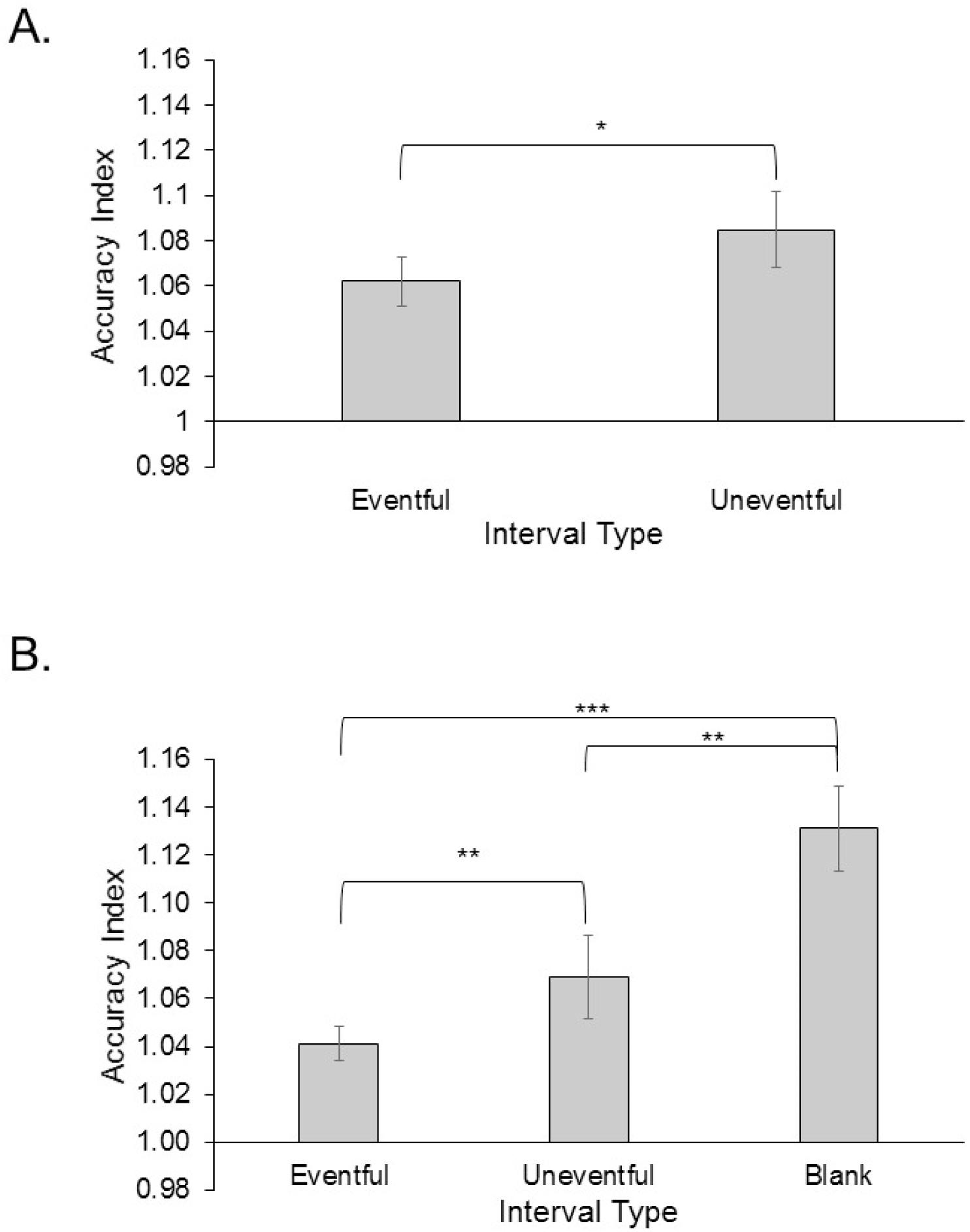
Panel A shows the AI values for eventful and uneventful movie intervals for Experiment 1. Panel B shows the AI values for eventful, uneventful and blank movie intervals for Experiment 2. Error bars are ± 1 standard error. *p < .05, **p < .01, ***p< .001

### Coefficient of variation (CV)

There was a significant difference between eventful (*M* = .19) and uneventful (*M* = .17) intervals, *t*(41) = 2.52, *p* = .016, *d* = .43. People made more variable reproductions during eventful movie intervals.

## Discussion

When individuals timed prospectively while watching movies of everyday events, the degree of eventfulness had an online influence on their temporal reproductions. In particular, participants made shorter reproductions during eventful than uneventful intervals, consistent with the possibility that more events led participants to believe more time had passed. Participants also showed more variable reproductions during eventful intervals, reflecting the larger range of eventfulness of these intervals compared to uneventful intervals. The fact that participants performed well on the order memory task suggests they attended to the movie content. These findings support EST and the view that automatic processing of online contextual changes during reproduction influenced prospective estimates of time even when the initially encoded interval involved no contextual changes.

If events had pulled attention away from the timing task, consistent with the AGT, individuals would have ended their reproductions later during eventful as compared to uneventful situations. However, it is notable that, overall, reproductions made while watching movies were longer compared to baseline trials. A possible explanation is that watching a movie draws attention away from timing, generally, but that within this timing context, eventfulness has an influence consistent with the expectations of EST. Experiment 2 sought to clarify the joint influence of attention and contextual change on prospective timing by comparing eventful and uneventful reproductions to a condition where participants simply watch a fixation.

### Experiment 2

For this experiment, we created “blank” movies for which individuals were cued to reproduce the target interval while watching a fixation with no eventfulness manipulation. If events serve as markers of time, then reproductions should be shorter for eventful than uneventful intervals, and the longest reproductions should occur during blank movies. If attending to movies, in general, pulls attention away from time, then blank movies should lead to the shortest reproductions.

## Method

### Participants

Sixty-six participants with normal vision and hearing completed this experiment for course credit. The experiment was approved by the Washington University Institutional Review Board. Four participants were excluded due to counting and one for failing to make reproductions on more than 33% of the experimental trials. This left 61 participants (mean age = 19.33 ± 1.04, 34 females) in the analyses.

### Apparatus/Stimuli

The same materials were used as in Experiment 1. However, we also included two “blank” movies for which participants reproduced the target while watching a fixation cross. These movies were created by identifying the initiation times of the intervals used for the 5 everyday movies and calculating the temporal distances between these time points. To create each blank movie, we randomly selected 5 distances out of the total to serve as the distances between each reproduction cue provided during these movies. After the final cue in each movie, we included a 50 s interval before the movie ended.

### Procedure

Participants completed the same temporal reproduction and order memory tasks and the same exit questionnaire as in Experiment 1; they also made reproductions during blank movies. The training procedure and instructions were identical except the blank movies did not involve an order memory task. Again, participants completed practice followed by the five experimental and two blank movies. Movie order was counterbalanced. Participants reproduced the target interval five times per blank movie to keep these movies roughly the same total length as the experimental movies. See Table 1 for the movie lengths.

### Data Analysis

The same trimming procedure from Experiment 1 was applied to experimental movie trials (both everyday and blank movies) and baseline trials and led to an average of .18 baseline trials and .41 experimental trials eliminated per participant. We calculated the AI for the baseline trials, eventful reproductions, uneventful reproductions and blank reproductions. CV was calculated for the different movie interval types.

Repeated measures ANOVAs were used to examine the impact of interval type (blank, eventful, or uneventful) on AI and CV. Where sphericity was violated, we report the Huyhn-Feldt correction. Main effects were investigated with post-hoc paired-samples t-tests, using the Bonferroni correction for multiple comparisons, which required a significance value of *p* < .017.

## Results

### Order Memory Task

Participants took on average 111.36 (*SD* = 34.06) seconds to complete the task and made an average of .56 (*SD* = .43) absolute deviations from the correct ordering of screenshots from the movies. This indicates that participants were attending to movie content.

### Accuracy Index (AI)

For baseline trials, participants achieved a mean AI of 1.03, indicating they achieved reasonable training on the target interval. We found a main effect of interval type, *F*(1.48, 89.01) = 18.22, *p* < .001, *MSE* = .010, *n_p_*^*2*^ = .23. People made shorter reproductions in eventful as compared to uneventful movie intervals, *t*(60) = 2.93, *p* = .005, *d* = .20, and compared to blank intervals, *t*(60) = 5.12, *p* < .001, *d* = .64. Participants also made shorter reproductions during uneventful relative to blank intervals, *t*(60) = 3.59, *p* = .001, *d* = .44 (see Figure 3).

### Coefficient of variation (CV)

There was a main effect of interval type, *F*(1.81, 108.31) = 5.45, *p* = .007, *MSE* = .003, *n_p_*^*2*^ = .08. Post-hoc t-tests indicated that people made more variable reproductions during eventful (*M* = .18) as compared to uneventful (*M* = .16) intervals, *t*(60) = 2.95, *p* = .005, *d* = .38 and compared to blank (*M* = .16) intervals, *t*(60) = 2.59, *p* = .012, *d* = .42. However, variability did not differ between uneventful and blank movie intervals, *t*(60) = .42, *p* = .675.

## Discussion

The findings from Experiment 2 replicated those of Experiment 1, with participants showing shorter and more variable reproductions during eventful relative to uneventful or blank movie intervals. This supports EST and runs counter to the expectations of the AGT. Participants performed well on the order memory task, suggesting they attended to the content of the movies. Interestingly, participants made the longest reproductions during blank movie intervals when there were no concurrent task features that would interfere with timing. If movies drew attention away from timing, the longest reproductions should have occurred during the movie interval reproductions. Instead, the data indicate that the most powerful impact on timing performance in the current experiment was the eventfulness of the experience in which participants made their temporal reproductions.

### General Discussion

In two experiments, we demonstrated that events in naturalistic stimuli influence prospective temporal reproductions in an online fashion. This is novel because most research showing event change effects on prospective timing have manipulated these changes at encoding and assessed participants’ subsequent reproductions or estimates (Boltz, 2005; Faber & Gennari, 2017; Waldum & Sahakyan, 2013), whereas we asked participants to reproduce a time duration during concurrent event processing during the reproduction phase. We demonstrated that participants were sensitive to the eventfulness of the elapsing interval and that more eventful movie intervals led them to think more time had passed, resulting in shorter reproductions. The current results support EST and the notion that events serve as temporal markers (Poynter, 1983).

Our findings do not support the AGT and, in fact, longer reproductions during blank movies than everyday movies runs counter to the idea that events draw attention away from timing. Instead, this result is consistent with blank movies proving to be the least eventful condition in the current study. Overall, these findings provide additional evidence that we automatically process event changes in our everyday experience and that doing so impacts our online sense of how much time is passing. For the caterer without a clock, if serving the main course was very eventful, she might think her staff was taking more time to complete this activity than they actually were, prompting her to hire more serving staff.

## Acknowledgments

We would like to thank Albert Deng, Elisa Kim, and Ed Bryant for their help with participant testing. The authors report no conflicts of interest. This work was supported by an NIA training grant: NIA Training Grant AG000030.

